# Genes on Different Strands Mark Boundaries Associated with Co-regulation Domains

**DOI:** 10.1101/2020.09.22.303438

**Authors:** Audrey Baguette, Steve Bilodeau, Guillaume Bourque

## Abstract

Gene regulation is influenced by chromatin conformation. Current models suggest that topologically associating domains (TADs) act as regulatory units, which could also include distinct co-expression domains (CODs) favouring correlated gene expression. We integrated publicly available RNA-seq, ChIP-seq and Hi-C data from A549 cells stimulated with the glucocorticoid dexamethasone to explore how differentially expressed genes are co-regulated among TADs and CODs. Interestingly, we found that gene position and orientation also impact co-regulation. Indeed, divergent and convergent pairs of genes we enriched at sub-TAD boundaries, forming distinct CODs. We also found that genes at COD boundaries were less likely to be separated by structural proteins such as Cohesin and CTCF. A complementary analysis of lung expression quantitative trait loci (eQTL) demonstrated that genes affected by the same variant were more likely to be found on the same strand while lacking a TAD boundary. Taken together, these results suggest a model where gene orientation can provide a boundary between CODs, at the sub-TAD level, thus affecting their likelihood of co-regulation.

## INTRODUCTION

There is mounting evidence that chromosome structures are important for gene regulation. Techniques such as genome-wide chromosome conformation capture (Hi-C) have tried to elucidate chromatin contacts in the 3-dimensional nuclear space (1–3). At the lowest level of resolution, compartments can be identified from Hi-C data. Compartments are divided in two types, A and B, defined by their interaction frequencies; compartments tend to interact more often with compartments of the same type (1, 4). Compartment A has been shown to contain more genes and to be enriched for active genes and open chromatin marks, while compartment B is enriched for closed chromatin marks (1, 5–7). Compartments are distinct from topologically associating domains (TADs), another structure found with high-resolution Hi-C. TADs are defined by their high concentration of contacts within their domain, relative to their low level of interaction across different TADs (8–11). Intra-TAD contacts vary from cell to cell (3, 12–14), but most TAD boundaries are conserved across single cells (3), cell types (7, 9, 11, 15) and species (9, 15, 16). In addition to TADs, that range from a few kilo bases to around 1 million bases long (8–11), smaller TAD-like structures, called sub-TADs have been discovered within TADs (4, 12–14). Finally, at the highest resolution, chromatin loops can be observed. They represent contacts between DNA regions, including promoters and enhancers (7, 13, 17–19), usually stabilized by CTCF or other structural proteins (15, 16, 20).

Several studies have confirmed that TADs construct an environment favouring gene co-regulation. Genes within the same TAD tend to be co-expressed (10, 16, 17, 21). Indeed, some TADs act as regulatory units after hormonal induction (22) and hormone responsive genes are found within the same interaction networks (23) or in-between TAD boundaries (24). Moreover, paralogs are usually co-regulated and found within the same TADs (25). However, there is also evidence that some genes resist the intra-TAD co-regulation (26). Co-regulation might thus be driven by sub-TAD structures such as cis-regulatory domains (27), insulation neighbourhoods (28) or co-expression domains (CODs) (26). Those seemingly contradictory observations show the shortcomings of our understanding of transcription with respect to chromatin architecture (8, 13). CODs are especially interesting to us as they are purely defined from a transcriptional point of view. Indeed, they are defined as regions of consecutive genes with correlated expression. It would thus be interesting to compare CODs to the well-known TADs and to better understand their similitudes and differences, in structure and in function. A few studies have found that divergent genes may share a promoter, so their expression is co-regulated (29, 30). However, the impact of gene orientation in relation to these higher-level structures (TADs and CODs) at the genomic level remains to be tested.

Here we propose a detailed characterization of the regulation boundaries at the TAD and sub-TAD level using a combination of available RNA-seq, ChIP-seq and Hi-C data from A549 cells induced with dexamethasone. We hypothesize that TADs and CODs have different types of boundaries. We looked at the properties of genes around these boundaries and found that the strand position of genes had a significant and differential impact. We further tested these observations using eQTL in lung cells. Our results lead us to propose a model for human cells where genes on the same stand have a high probability of co-regulation. COD boundaries, marked by the change of strand, reduce the co-regulation probability independently of structural proteins.

## MATERIAL AND METHODS

### Data origins and pre-processing

The data sets used in this paper come from previous studies on A549 cells and are available in public databases. The cells were all treated with 100nM of dexamethasone (DEX) or only a vehicle (for controls). RNA-seq (ENCSR897XFT), ChIP-seq (ENCSR571KWZ, ENCSR375BQN, ENCSR588JLN, ENCSR210PYP, ENCSR022IHB, ENCSR625DZB, ENCSR738NGQ, ENCSR447VJR, ENCSR180FFI, ENCSR868FCL, ENCSR342NKR, ENCSR476OXC, ENCSR790OOG, ENCSR483SDK, ENCSR501UJL, ENCSR376GQA) and Hi-C samples (ENCSR842RTB, ENCSR435JUA) were produced by the same laboratory (Dr. Tim Reddy, Duke) and the pre-processed files, annotated with the GRCh38 reference assembly, were retrieved from ENCODE (https://www.encodeproject.org/) (23, 24, 31, 32) (Supplementary tables 1-4).

For polyA^+^ RNA-seq, the trimmed, aligned and quantified files containing the raw read counts for each gene were downloaded, for an hourly time-course (Control, 30m, 1h, 2h, 3h, 4h, 5h, 6h, 7h, 8h, 10 and 12h), each timepoint having three or four replicates. Genes are labeled with their ENSEMBL name and read counts for the multiple isoforms were summed. The most upstream base of all isoforms was selected as the start of the gene and the most downstream as the end. Quality control was made by normalizing the read counts with edgeR (https://bioconductor.org/packages/release/bioc/html/edgeR.html) (33, 34) and removing batch effects with svaseq (from the sva package, https://bioconductor.org/packages/release/bioc/html/sva.html) (35) following the method used by McDowell et al. (24). A principal components analysis (PCA) analysis and a t-distributed stochastic neighbor embedding (t-SNE) analysis were performed on normalized read counts. The replicates clustered correctly by timepoint.

For ChIP-seq data sets, trimmed and aligned *bam* files and *bed* files resulting from peak calling were downloaded for all available replicates, conditions and targets. The ChIP-seq datasets follow the same hourly time-course as the RNA-seq data set. A consensus peakset was produced using DiffBind (https://bioconductor.org/packages/release/bioc/html/DiffBind.html) (36, 37). DiffBind uses the peaks called at each timepoint and the sequenced reads to create RNA-seq-like read counts, with the number of reads found in each replicate, at each timepoint, for all called peaks. We were not interested in performing a differential binding analysis, so the read counts were not considered. We only used the consensus peakset produced by DiffBind to identify binding sites across the time-course.

Concerning Hi-C data, in addition to the *hic* files available for all replicates, the already identified topologically associated domains contained in *bedpe* files were also downloaded. The Hi-C time-course contains only 5 points: 0h, 1h, 4h, 8h and 12h. The changes in chromatin were quantified using 3DChromatin_ReplicateQC (http://github.com/kundajelab/3DChromatin_ReplicateQC) (38), which itself implements four different quality control methods for Hi-C data. First, the pre-processed *hic* files were downloaded from encode and dumped using the *dump observed* method of Juicer (39) using bins of 10kb. That resolution was selected as it was the middle resolution the three resolutions used to call TADs (5kb, 10kb and 25kb) (23). 3DChromatin_ReplicateQC first computes quality scores using QuASAR-QC (40), then it uses QuASAR-Rep (40), GenomeDISCO (41) and HiC-Spector (42) to produce reproducibility scores. QuASAR uses transformed matrices, based on read count matrices and enrichment matrices, corrected for distance, to produce quality scores for all chromosomes and replicates.

TAD boundaries were created by directly taking the list of found TADs at the time 0 available on ENCODE and creating regions 500 bp upstream and 500 downstream of all beginning and ends of TADs. Overlapping boundary regions were merged, such that the 5935 TADs resulted in 11454 TAD boundaries.

### Categorizing and pairing genes

First, differentially expressed genes were identified individually for each timepoints using edgeR and DESeq2 (https://bioconductor.org/packages/release/bioc/html/DESeq2.html) (43). A gene is identified as upregulated if it is differentially expressed according to both methods (FDR < 0.05 for edgeR and padj < 0.05 for DESeq2) and that have a higher expression level at the considered timepoint than the reference. Downregulated genes are those which are differentially expressed and have a higher expression level in the controls. A consensus was created across the time-course such that “Up” genes are genes that are found to be upregulated for at least two timepoints but are never downregulated. The same principle is applied to identify “Down” genes. Genes that did not fall in either of those categories were said to be “Stable”. All genes with detectable expression were considered. The “Stable” category contains slightly more genes with low expression levels, but the overall distribution of the mean expression of genes is similar for “Stable”, “Up” and “Down” genes (Supplementary figure 1).

Genes were then paired, and their distance was calculated, from TSS to TSS. Only pairs of genes separated by less than 1Mb, from TSS to TSS were kept. This resulted in six categories of pairs, depending on their comportment: “Up-Up”, “Down-Down”, “Stable-Up”, “Down-Stable” and “Down-Up”.

With the 14493 genes having detectable transcription, 147248 pairs separated by less than 1Mb could be formed. In addition, pairs were labeled according to the relative position of the genes: “Same strand” if both genes are located on the same DNA strand, “Divergent” if the genes are on different strands and back-to-back and “Convergent” if they are on different strands and facing each other.

### Odds ratios and distribution matching

As one of the main objectives is to find whether genes with opposite behaviours are separated by a physical barrier, the enrichment for finding a barrier between such pairs had to be computed. Enrichments were expressed in odds ratios, where the “Stable-Stable” category was used as reference. However, the distribution of distances is not the same between pair types. To avoid a bias where pairs separated by a larger distance are found to have a barrier between them by chance, the reference pairs were sub-sampled such that the distribution of distances of the resampling matches the distribution of distances of the pairs of interest, and the query and reference contain the same number of pairs. The distribution matching algorithm consists in dividing the distributions of distance from the interest pairs and the control pairs into bins of 5kb, then counting the number of interest pairs falling in each bin and sampling as many control pairs in the corresponding bin. The resampling was done 1000 times, to allow the calculation of empiric p-values. Odds ratios are defined as follows:

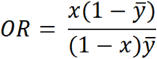

In the above equation, *x* corresponds to the proportion of pairs of interest containing a barrier between the two genes and *y* represents the proportion of pairs of interest, after resampling, with a barrier. The region “between the two genes” is considered as the regions between gene bodies, thus overlapping genes cannot contain a barrier between them, even if their TSS are not at the same position. For this analysis and the following, only the pairs with genes separated by less than 100kb were kept, which resulted in 20197 pairs.

### Characterizing boundaries

To characterize the boundaries, only the pairs of consecutives genes were kept, which totalize 8946 pairs of genes. The pairs of genes were subdivided in categories depending on whether they were found in the same TAD, around a TAD boundary or outside TADs. Pairs within the same TAD were further subdivided between pairs where both genes have the same behaviour (“Up-Up”, “Down-Down” and “Stable-Stable”) or opposite behaviour (“Down-Up”, “Down-Stable” and “Stable-Up”). The relative abundance of same-strand, convergent and divergent pairs was compared for those four categories: 1-pairs at TAD boundary, 2-pairs outside TADs, 3-pairs inside the same TAD with the same behaviour and 4-pairs inside the same TAD with opposite behaviours. That last category serves as a proxy to COD boundaries. The pairs were then analysed separately depending on the “strandedness” to find the repartition of structural proteins such as CTCF and Cohesin. As Cohesin does not have direct ChIP-seq data, the co-localization of RAD21 and SMC3, two of its subunits for which there are available ChIP-seq replicates, was used instead.

The odds ratio heatmap has been by comparing convergent and divergent pairs to “same-strand” pairs after consecutive resampling to match distance distributions as before. Unlike in the first heatmap, the empirical p-values were computed considering both tails, rather than just the upper tail, to be able to have p-values associated with depletion too and not only with enrichment.

### Pairing eQTL targets

The significant eQTL and their gene targets found in healthy lung cells were retrieved from GTEx (https://gtexportal.org/home/) (44). The data used for the analyses described in this manuscript were obtained from Single-Tissue cis-eQTL Data on the GTEx Portal, dbGaP accession number phs000424.v8.p2 on 03/02/2020. As we wanted to analyse the relation between genes affected by the same eQTL, all eQTLs with a single target were discarded. This resulted in a total of 477420 selected eQTLs and 9434 gene targets. All genes having their TSS within 100 kb of each other were paired, but pairs involving the same genes were only considered once. Indeed, two genes can be both affected by multiple eQTLs and we wanted to consider each pair of genes uniquely, even if multiple variants are involved. We thus obtained 9088 interest pairs, where both genes are affected by the same eQTL, and 24035 control pairs, where genes are affected by different eQTLs or one gene is affected by an eQTL and the other is not. The prevalence of finding those genes on the same strand or not, or to be separated by a barrier or not was assessed following the distribution matching technique and the odds ratio formula described previously. Two genes were said to be separated by a barrier if there was either a TAD boundary or if there was evidence of CTCF, RAD21 and SMC3 between them. The pairs of genes affected by the same eQTL were compared to pairs in which one gene is affected by an eQTL and the other is not significatively impacted by it and is within 100kb of the first TSS. To account for the variations in which eQTL affect genes, we tested all pairs in addition to the following subsets: 1-only pairs of consecutive genes, 2-all pairs affected by strong eQTLs (regression slope of the gene-eQTL association > 0.7 or < −0.7) and 3-consecutive pairs affected by strong eQTLs. The pairs were then further categorized into 1-pairs on the same strand without barrier (both CTCF and Cohesin or a TAD boundary) between them, 2-pairs on opposite strands without barrier between them, 3-pairs on the same strand with a barrier between them and 4-pairs on opposite strands with a barrier between them. P-values associated with both depletion and enrichment were computed empirically as before.

## RESULTS

### Gene expression changes associated with glucocorticoid stimulation

To study the relation between gene regulation, chromosome architecture and gene position/orientation, we integrated RNA-seq, ChIP-seq and Hi-C ENCODE datasets from A549 cells induced with 100nM dexamethasone (DEX) (31, 32). First, we reanalysed the RNA-seq data to define differentially expressed genes following DEX stimulation (Material and Methods). Distribution of the RNA-seq datasets was consistent with progressive changes in gene expression with time (Figure 1A). This observation was supported by a principal component analysis showing the same progression following the first and second principal components (Supplementary figure 2A). Next, differentially expressed genes identified using edgeR (33, 34) and DESeq2 (43) were retrieved at each timepoint. To define a consensus set of differentially expressed genes, we selected genes that were differentially expressed at least twice (in the same direction) during the time-course. This provided us with a list of 1716 upregulated genes (labelled as “Up”) and 1810 downregulated genes (labelled as “Down”). Furthermore, a total of 10751 were unchanged (labelled as “Stable”) (Supplementary table 5).

**Figure 1.**
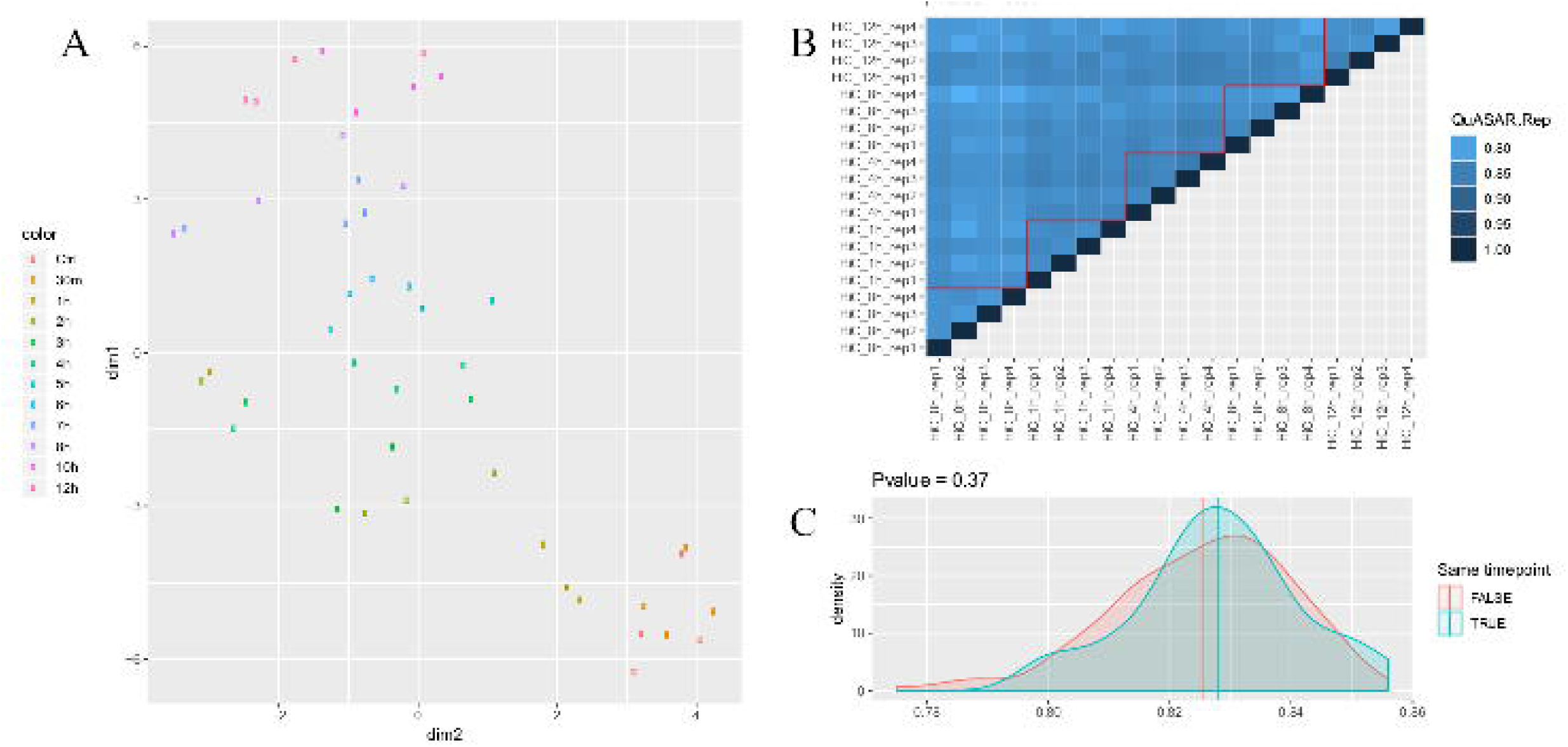
Gene expression changes in the RNA-seq samples and Hi-C reproducibility scores. (A) t-SNE of the normalized RNA-seq data. (B) Heatmap of the replication scores given by QuASAR-Rep. Comparaisons inside the same timepoint are under the red lines and comparisons across timepoints are over it. (C) Distributions of the scores between replicates of the same timepoint (blue) or different timepoints (red). The vertical lines represent the means of the distributions. The p-value comes from a t-test.

### The overall chromosome architecture is maintained during DEX stimulation

To determine the chromosome architecture of the A549 cells during DEX stimulation, including TADs, we used Hi-C datasets at 0h, 1h, 4h, 8h and 12h (Material and Methods). We assessed the quality of these Hi-C datasets using QuASAR-QC (40). The quality scores for all chromosomes were found to vary between 0.015 and 0.02 (Supplementary figure 2B), which is characteristic of somewhat noisy data at this resolution of 10kb, but not uncommon (38, 40). The reproducibility between pairs of replicates, from the same timepoints and across timepoints, was then compared using three methods: QuASAR-Rep (40), GenomeDISCO (41) and HiC-Spector (42). The reproducibility scores were used to quantify the similarity of the Hi-C maps through the time-course. Indeed, Hi-C maps and the TADs derived from them were available for five timepoints and we wanted to see if the maps are similar enough to only use a single timepoint as reference to locate TAD boundaries. The reproducibility scores varied from method to method, but all methods confirm that there is no more variability between replicates from different timepoints than between replicates from the same timepoint (Figure 1B, Supplementary figure 2C and E). This can be visually assessed with the heatmaps of the scores and is confirmed by a two-sample t-test comparing the distributions of the pairwise scores (Figure 1C, Supplementary figure 2D and F). None of the p-values for QuASAR-Rep (p-value = 0.37), GenomeDISCO (p-value = 0.36) and HiC-Spector (p-value = 0.22), respectively, was found to be significant. We cannot draw conclusions as to whether the TADs are completely stable after DEX induction using only those scores. However, as the scores report no major change in chromatin architecture, they justify the use of a single timepoint as reference for TAD boundaries to facilitate the analyses below.

### Co-expressed genes show genomic proximity

We paired all genes that were previously characterized (“Up”, “Down” and “Stable”) to create a compounded matrix of pairs of genes. Then, we kept pairs for which the distance between the transcription start sites (TSS) was less than 1 Mb (Material and Methods), providing 147,248 pairs. Indeed, pairs separated by less than 100kb make up 13.72% of all pairs, but as much as 19.08% of pairs showed concurrent upregulation compared to only 6.8% of pairs where one gene is activated and the other is repressed (Figure 2A). This is seen at a finer scale too, as even when only the pairs separated by less than 100 kb are considered, the distributions of distances changes depending on the relative behaviours of the genes within the pair (Figure 2B). On average, activated pairs were separated by 14651 bp, while 73207 bp for genes going in opposite directions. There is thus a distance bias to consider when comparing the elements separating different types of pairs.

**Figure 2.**
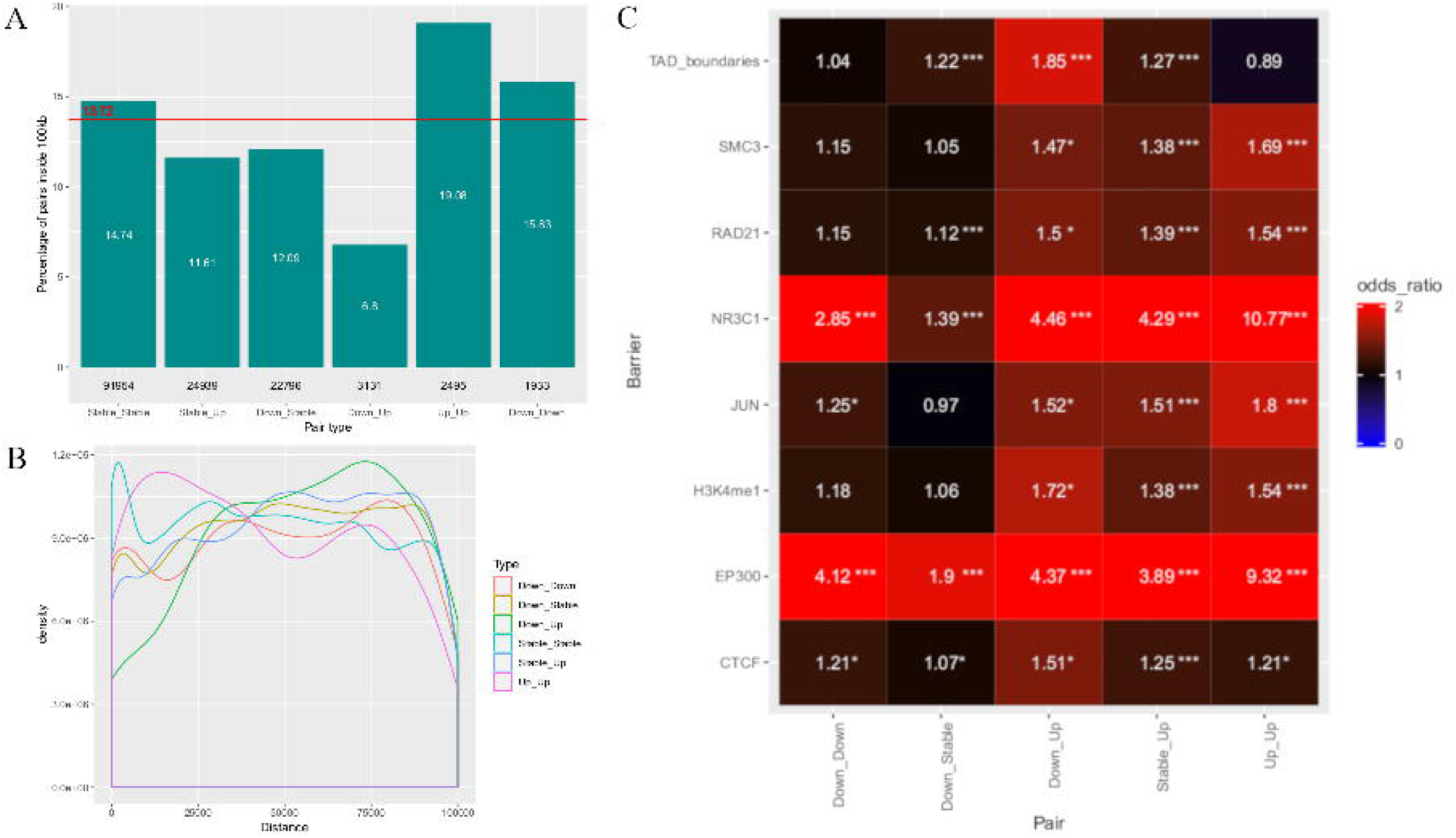
Genes with the same behavior tend to be closer from each other and pairs of genes going in opposite directions are more often separated by TAD boundaries than pairs of stable genes. (A) Proportion of pairs in which genes are separated by less than 100kb among all pairs in which genes are separated by less than 1Mb. The red line represents the proportion when all pairs are considered. (B) Distribution of distance (in bp) between the genes of the pairs, for pairs separated by less than 100kb. (C) Heatmap of odds ratios for the presence of a physical barrier between the genes of the pairs. The “Stable_Stable” pairs are used as reference. *P-value < 0.05; ***P-value < 0.001. The p-values are empirical, computed with 1000 resampling events.

TAD boundaries are known to have insulator properties, limiting the effects of enhancers to nearby genes on the other side of the boundary (9, 12, 14, 45). If true, the prevalence of TAD boundaries between genes having opposite behaviours following DEX induction should be higher. We retrieved the list of TADs, called with Juicer’s Arrowhead (39), at the earliest time-point (0h) (23, 31, 32). We then transformed the TADs list in a list of 11454 TAD boundaries (Material and Methods, Supplementary figure 3). Stable gene pairs were used as a reference throughout. To account for the identified distance biases, the reference pairs were sub-sampled 1,000 times, such that the distribution of distances of the sub-sampling matches the distribution of distances of the pairs of interest and the query and reference contain the same number of pairs (Supplementary figure 4). As expected, TAD boundaries were found to be significantly enriched between pairs of genes having a different behaviour, especially pairs where one gene is activated and the other is repressed (odds ratio of 1.85, empirical p-value < 0.001) (Figure 2C, Supplementary figure 5A). This enrichment is also seen when comparing pairs of genes with different behaviour to pairs with genes having the same behaviour, suggesting the enrichment might have a real biological significance and is not only an artefact due to the low level of expression of “Stable” genes relative to “Up” and “Down” genes (Supplementary figure 5B). On the contrary, pairs with genes moving in opposite directions, TAD boundaries were not enriched (odds ratio of 1.04 and 0.89, p-values > 0.05).

#### CODs are not delimited by TAD boundaries

While TADs represent a physical structure part of the chromatin, CODs are defined as chromosome regions containing a number of genes being co-expressed. To determine the nature of the boundary separating CODs, we analysed the chromatin features between each pairs of genes. Indeed, while positive and significant, the previous odds ratios suggest that CODs are not primarily separated by a TAD boundary. If TAD boundaries are not a structural determinant separating CODs, we wondered if other chromatin features could be found. We thus looked for factors which presence would be enriched between pairs of genes with an opposite behaviour. The available raw reads and pre-called peaks of the ChIP-seq data corresponding to the dataset (24) were further processed through DiffBind (36, 37) in order to find the binding sites of the 16 proteins along the time-course (Material and Methods). This resulted in, for example, 52729 CTCF peaks, 14699 NR3C1 peaks, 72433 RAD21 peaks and 71220 SMC3 peaks (Supplementary figure 3).

The enrichment between genes with opposite behaviours is weaker for CTCF (odds ratio of 1.51, p-value < 0.05) (Figure 2C). In addition, pairs of upregulated genes and pairs of downregulated genes do not show any enrichment for TAD boundaries but only small, significant enrichments for CTCF (odds ratios of 1.21 in both cases), confirming CTCF alone is not enough to create the insulation property of TAD boundaries. SMC3 and RAD21 are enriched around activated genes (odds ratio of 1.69 and 1.54, respectively, empirical p-value < 0.001), which is consistent with their function; those are two subunits of Cohesin, a ring-shaped protein complex that helps bringing promoters and enhancers in close proximity (15, 16, 20). NR3C1 is directly activated by DEX and is thus expected to be located around differentially expressed genes, especially activated genes, and this is confirmed by its enrichment pattern across the gene pairs. NR3C1 is also very significantly enriched between pairs of downregulated genes, confirming GR acts as a repressor and not only as an activator. As EP300 binds to enhancers, it is found around expressed genes. Indeed, we see an enrichment of EP300 between genes that were highly expressed at the start of the time-course (downregulated genes) or at the end (upregulated genes). Taken together, these results show that our distance-bias correction permits to compare the proportion of proteins found in-between pairs of genes, but that none of the proteins were found to form COD boundaries.

#### Gene conformation marks sub-TAD boundaries

Since we were unsuccessful in finding factors constitutively present at COD boundaries, we turned our attention toward the genome structure at a smaller scale. Therefore, we decided to measure “strandedness” (convergent, divergent or same strand) in regard to the presence of COD boundaries. To better differentiate TAD boundaries from COD boundaries, four new categories of gene pairs were created, depending on whether they were 1-around a TAD boundary, 2-inside the same TAD but had different behaviours (potentially associated with a COD boundary), 3-inside the same TAD but with the same behaviour (thus probably within the same COD), or 4-outside TADs (Material and Methods). To characterize the relation of genes at the boundaries, only the pairs of genes that were the closest to the boundaries were considered. Analysing the distribution of pairs in different conformations according to those categories showed that at TAD boundaries, gene pairs are usually separated by both Cohesin (SMC3 and RAD21) and CTCF, regardless of the relative position of genes (Figure 3A). However, while inside TADs, divergent and convergent pairs of genes are usually less often separated by CTCF and Cohesin than same-strand pairs. Indeed, among pairs of genes found within the same TAD that show a different behavior (“Down-Stable”, “Down-Up” and “Stable-Up”), 64.6% of same-strand pairs have CTCF and Cohesin between them, while it is only true for 50.5% of the convergent and 42.2% of divergent pairs. If we consider pairs of genes found in the same TAD that also have the same behavior (“Down-Down”, “Stable-Stable” and “Up-Up”), we find that 50.8% of the same-strand pairs have both structural proteins, against as few as 42.1% for convergent pairs and 31.1% for divergent pairs. Using the “same strand” pairs as reference, while accounting for any possible distance bias (Material and Methods), divergent and convergent genes were found to be significantly depleted of CTCF and Cohesin everywhere (empirical p-value < 0.001) but at TAD boundaries as compared to same strand pairs with a same distribution of distances between the genes (Figure 3B). The odds ratios for finding CTCF and Cohesin between divergent genes are of 0.52 when genes are in the same TAD but have a different behavior, of 0.64 when genes are in the same TAD and show the same behavior and of 0.58 when genes are outside TADs (empirical p-values < 0.001). For convergent genes, the odds ratios are of 0.61, 0.75 and 0.56 (empirical p-values < 0.001), in the same order. Assuming pairs of genes with a different behavior mark COD boundaries, the depletion of CTCF and Cohesin between divergent and convergent pairs suggests that those pairs could mark sub-TAD domains without the need of Cohesin or CTCF.

**Figure 3.**
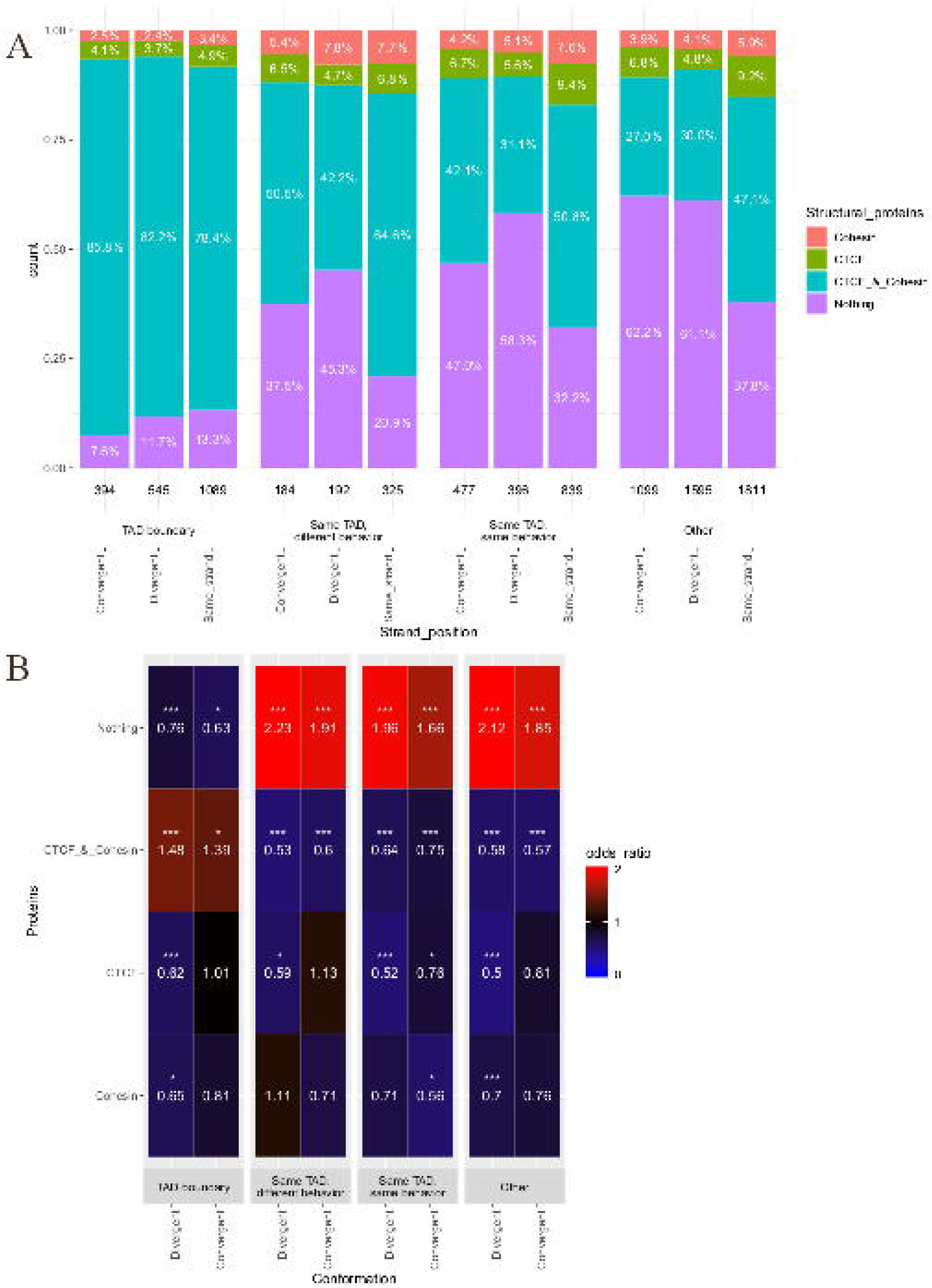
Repartition of the pairs of consecutive genes and of structural proteins across boundaries. (A) Proportion of convergent, divergent and same-strand pairs of consecutive genes having structural proteins between them in each location category. (B) Heatmap of the odds ratios, as compared to the “same strand” pairs in the same category. *P-value < 0.025 or p-value > 0.975; ***P-value < 0.001 or p-value = 1. The p-values are empirical, computed with 1000 resampling events.

#### eQTL gene targets are preferentially on the same strand

Our results support the idea that CODs are often separated by genes found on opposite DNA strands. If this is true, we hypothesized that expression quantitative traits loci (eQTL) affecting multiple genes will also be found to preferentially regulate genes on the same DNA strand. To test this model, we collected eQTL for lung tissue to match A549 cells from the GTEx Portal (44). We compared 9088 pairs of genes affected by the same eQTL to 24035 control pairs using the same resampling technique to limit the distance bias as before (Material and Methods). We found that pairs of genes affected by the same eQTL show a preference for being on the same strand, without a strong barrier such as a TAD boundary or the co-localization of CTCF and Cohesin (Figure 4A). Indeed, 19% of pairs fall under that category, while the expected proportion is of around 14% for pairs of genes not affected by the same eQTL (odds ratio of 1.44, empirical p-value < 0.001). The difference is even more marked when pairs of genes affected by the strongest variants are considered. Indeed, the proportion of pairs affected by the same genes that are found on the same strand without barrier between them goes as high as 33%, resulting in an odds ratio of 2.68 (empirical p-value < 0.001, Figure 4B). Moreover, the pairs are less likely to be on opposite strands and separated by a barrier than control pairs. 58% of all pairs of interest fall under that category, while the expected value is around 66% (odds ratio of 0.73, empirical p-value < 0.001). Once again, considering the most affected genes makes the contrast even stronger, as the proportions drops to 39% and the odds ratio to 0.4 (empirical p-value < 0.0001). To confirm that the definition of the space “between genes” did not influence the results, we reanalysed the data using the intergenic space instead of the space between the TSS which confirmed the main conclusions (Supplementary figure 6). Specific examples of those observations include the eQTLs chr1_109678559_T_A_b38 and chr3_195614883_T_A_b38 (Figure 4C-D). Taken together, our results support that eQTLs affect preferentially genes located on the same strand and their action is limited by barriers, including opposite strand gene conformation.

**Figure 4.**
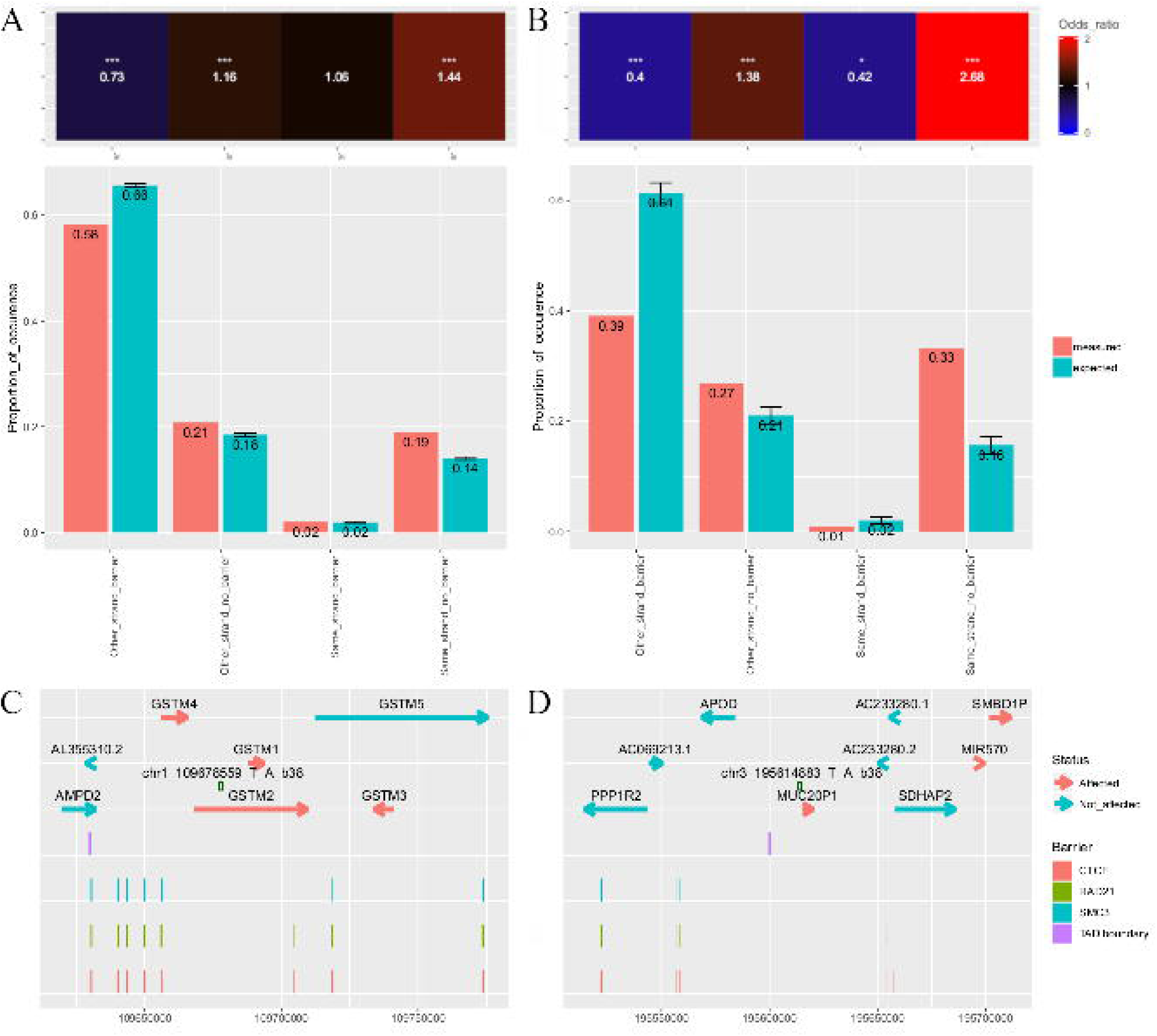
Analyzing gene conformation with eQTLs. Pairs of genes where both genes are affected by the same eQTL are enriched for being on the same strand without barrier (CTCF and Cohesin or a TAD boundary) and depleted for being on different strands and separated by a barrier, as compared to control pairs. This is true when considering (A) all pairs of genes separated by 100kb affected by the same eQTL or (B) a subset with only the pairs with the strongest association (regression slope > 0.7 or < −0.7) to their eQTL. *P-value < 0.025 or p-value > 0.975; ***P-value < 0.001 or p-value = 1. The p-values are empirical, computed with 1000 resampling events. (C-D) Specific examples of variants (green dots) and the genes they affect depending on their orientation and the position of barriers.

### DISCUSSION

In this study, we compared the insulation property of TAD and COD boundaries. Genes having an opposite behavior after DEX induction are enriched to have a TAD boundary between them, with an odds ratio of 1.85 as compared to pairs of stable genes. The odds ratio is smaller (1.51) for having CTCF between them. We also show that the relative position of genes seems to create structural protein-independent boundaries inside TADs. At TAD boundaries, same-strand, convergent and divergent gene pairs are usually separated by CTCF and Cohesin. However, when pairs are inside the same TAD, more than half of the same-strand pairs still contain CTCF and Cohesin, but the convergent and divergent pairs are not separated by those proteins as often. Finally, we further supported the observations using eQTL data and found that genes affected by the same eQTL are enriched for being found on the same strand, without barrier, than genes not affected by the same variant (odds ratio of 1.44). Moreover, genes affected by the same eQTL are depleted for being on different strands and separated by a barrier (odds ratios 0.73). When the genes most affected by eQTLs are considered, the tendencies are even more marked. Taken together, all those results suggest that convergent and divergent pairs mark the boundaries between co-expression domains, at the sub-TAD level, without the need of CTCF and Cohesin.

#### A new, probabilistic model for gene co-regulation

Our observations suggested a model for human cells where consecutive same-strand genes make up small sub-TAD domains within which genes are likely to be co-expressed (Figure 5A). The boundaries of those domains are marked by the change of strand, as genes on different strands are less likely to be co-expressed. The cell can introduce CTCF and Cohesin that tend to create insulation boundaries such as TAD boundaries that disrupt the possibility of co-regulation (Figure 5B). Those small sub-TAD domains are the building blocks of co-regulation domains. CODs are, by definition, statistical entities and not physical entities, as they are found by aggregating all genes that have correlated expression. We propose that sub-TAD boundaries which are protein-independent mark the preferential split point of CODs. In a specific condition, genes on either side of the boundary can have similar expression level and make up a single COD. After a stimulus, the expression levels of the two regions might not be the same anymore and they would split into different CODs according to the position of the boundary (Figure 5C). The proposed model is not mutually exclusive with previous ones, such as transcription factories, and could help to understand how genes that do not seemingly receive the same signal could have similar expression levels. The relation between TADs and transcription is a subject that many studies tried to understand(10, 22, 26, 27). The boundaries of co-expression domains were not fully characterized and TADs were lacking a functional definition (13). Using publicly available data of A549 cells induced with dexamethasone (23, 24, 31, 32), we found that the strand position of genes influences the probability of their co-regulation and that TAD boundaries disrupt co-regulation.

**Figure 5.**
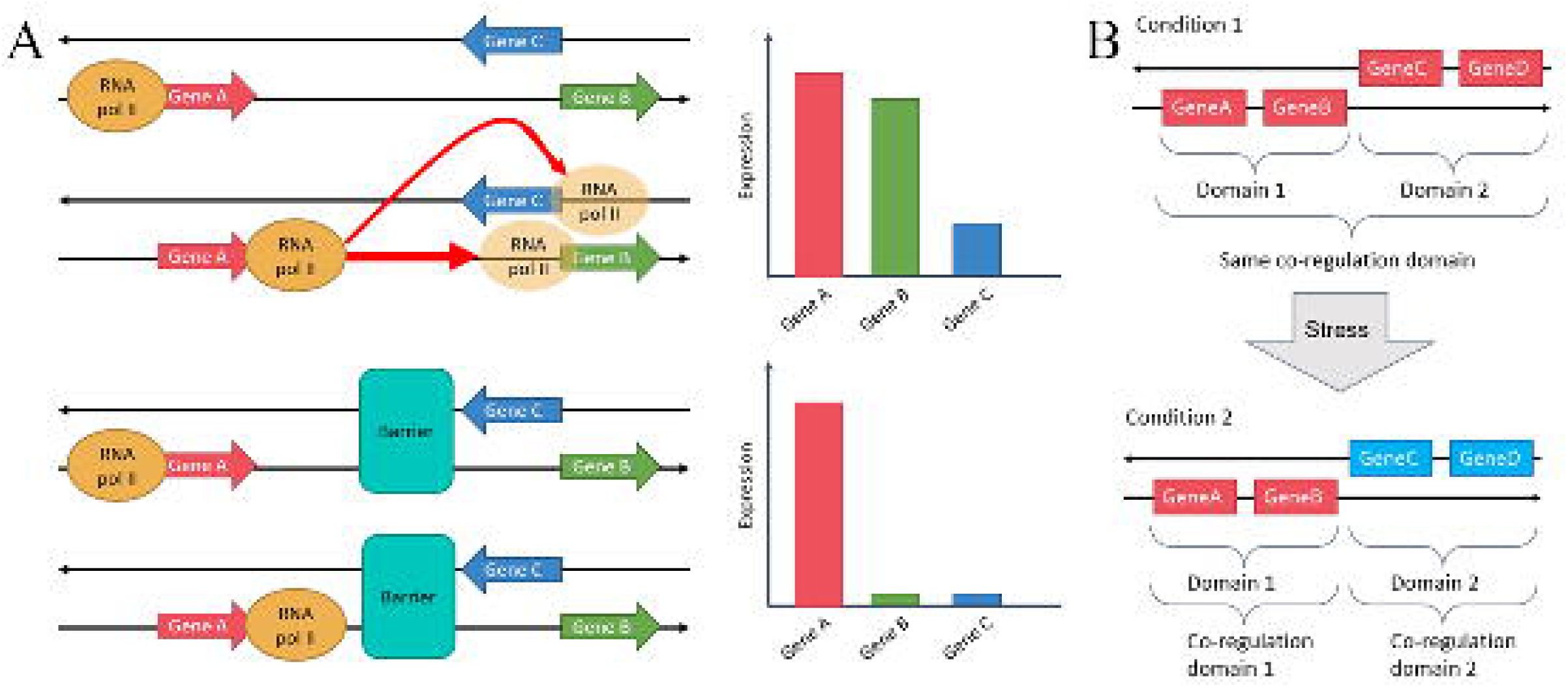
Model of co-regulation likeliness. (A) Same-strand genes are very likely to be co-expressed, as the RNA pol II just needs to continue its path along the strand. Divergent and convergent genes are less likely to be co-expressed, as the RNA pol II would needs to detach and reattach itself to go from one gene to the other. (B) When genes are separated by a barrier (CTCF and Cohesin or TAD boundary), there is complete disruption of co-regulation. (C) Illustration of how domains - physical entities - can make up a single co-regulation domain - a statistical entity - or split up in two, depending on the condition.

### Limitations and perspectives

The proposed model has only been observed in A549 cells and is thus limited to that cell line for now. It would be interesting to test it further using other conditions, in other cell lines or even different organisms. In addition, eQTL data comes from normal lung cells of over 850 individuals, while A549 cells come from a lung cancer cell line. Cancer cells usually contain structural variants which could impact the genomic architecture. The position of TAD boundaries, CTCF, RAD21 and SMC3 used during validation with eQTL comes from A549 cells. Those elements have been reported to be greatly conserved, but there still could remain differences that could influence our validation step. Ideally, for future validation of the model, Hi-C, ChIP-seq and eQTL data should come from the same cells. Using different cell types would also help COD boundaries detection. We used consecutive genes with a different behavior as proxy to COD boundaries. However, two genes could be found to be upregulated after DEX induction, but it does not mean they have completely correlated expression. They might thus be found to be into different CODs while our method would not identify it as boundary. We may thus potentially be missing COD boundaries. CODs also change depending on the cell needs and therefore, to have a complete mapping of CODs and their relative splitting or merging from one condition to another would be useful. Still, the fact that we can support our model with significant values knowing our definition of COD boundary is more stringent reinforces our confidence in the model.

An important control for the analyses presented in this study was to check for properties that could be different between the tested pairs of genes and the control sets that would introduce a bias. Looking at the distribution of distances between pairs of genes depending on their relative behaviour showed that there is a strong distance bias. This was accounted for but still consists the main limitation of the study, as it is reduced but cannot be completely removed. A second technical limitation is that the effect of TAD boundaries on co-regulation has been assessed using TAD boundaries derived from the TADs called before DEX induction (time 0). We used the reproducibility scores of the Hi-C maps to justify that selection. However, all possible changes in structure are then contained within a single number. Since the regions covered by TAD boundaries add up to a small portion of the genome, it is possible that small changes in TAD boundaries location would be lost in the general score given for the whole genome. It might thus be better to account for those small changes in future explorations of the model. Lastly, we assumed all genes have their own promoter. However, there are a few reported case of divergent genes sharing a promoter, that thus have coordinated expression (29, 30, 46). Those remain exceptional cases, but it would be interesting to treat genes sharing a promoter separately from all others when further exploring the model, as they constitute another mechanism for controlling co-regulation.

Despite the limitations of our method, the model for human cells we propose is supported by different types of analyses, using different data types, which leads us to believe in its robustness. Exploring it further with improved bias correction and supplementary data is more likely to improve the results than disprove them. Our model is also compatible with other suggested models. One rising paradigm change in how active transcription is seen suggests that RNA polymerase II is not the moving part during transcription, but rather that it is fixed in the nuclear space (47, 48). Multiple RNA polymerase II would cluster into transcription factories (47, 49–51) and DNA would cluster to factories, or detached from them, depending on the cellular needs (52). Our model has been explained such that RNA polymerase II is the mobile part, for ease of understanding, but it is not mutually exclusive with the idea of fixed polymerase. If the transcription machinery is indeed fixed, the model stays the same and can be explained as such: when two genes are located on the same strand, they are very likely to be co-expressed as the DNA could not detach itself from the transcription factory. When genes are divergent or convergent, they are slightly less likely to be co-expressed as the DNA would have to detach then re-attach itself to the factory, but slightly shifted to attain the TSS on the other strand. CTCF and TAD boundaries greatly decrease the probability of co-regulation as they serve as barrier, preventing the DNA to slide freely. The model we propose is thus compatible with the other models of transcription factories and might even help understanding how transcription is regulated in those hubs of active transcription. As such, it would serve as a basis for more complex gene regulation mechanisms and could well be the key to understand unresolved biological questions: What is the regulatory function of the genomic architecture (8)? What is the functional definition of TADs and sub-TAD domains (13)? How would cells lacking structural proteins behave following a stimulus demanding a change in the expression program (53)? We suppose that the regulatory function of the chromatin organization differs at depending on the level considered; gene “strandedness” affects the probability of co-regulation while TAD function is to disrupt the spread of expression signals through its boundaries. The functional definition is TADs would thus be centered on the insulation properties of the boundaries, while the sub-TAD domains definition would rather relate to gene position. Cells that do not have the necessary structural proteins would have expression patterns reflecting the relative position of the genes, with close, same-strand genes behaving similarly. The model thus serves as a stable ground on which complex hypotheses can be constructed and tested in the near future.

In this paper, we describe the existence of intra-TAD boundaries, delimited by the changing of strand on which genes are placed, that change the probability of co-regulation. The regions bordered by those new boundaries act as the building block of co-expression domains (COD). Indeed, CODs are statistical entities, created by clustering all consecutive genes having a correlated expression. However, correlation is different from causality and genes could have correlated expression by chance. If two nearby genes need to be expressed in similar amounts in condition A, they would be part of the same COD. If the cell enters condition B, the two genes might not be transcribed in similar amounts anymore and they would be split up into different CODs. There are thus regions bordered by physical boundaries (the change of strand), independent of structural proteins, within which genes have a probability of co-regulation, and that can be “assembled” to form CODs. To completely disrupt the possibility of having co-regulation of nearby genes, TAD boundaries or the co-localization of CTCF and Cohesin are introduced. Such model was validated using eQTL data, but further work would be needed to exactly determine if the model can be extended to other human cells or other organisms.

## AVAILABILITY

All R packages used are available on Bioconductor (https://www.bioconductor.org/). 3DChromatin_ReplicateQC, the algorithm used to compare Hi-C maps, is available on GitHub (http://github.com/kundajelab/3DChromatin_ReplicateQC).

## ACCESSION NUMBERS

All data has been retrieved from ENCODE (31, 32) and GTEx (44). ENCODE accession numbers are detailed in Supplementary table 1 to 4. GTEx data accession number is phs000424.v8.p2.

## SUPPLEMENTARY DATA

Supplementary Data are available at NAR online.

## ACKNOWLEDGEMENT

We would like to thank Rola Dali and Jacek Majewski for advice throughput the study and comments on the manuscript. Data analyses were enabled by compute and storage resources provided by Compute Canada and Calcul Quebec.

## FUNDING

This work was supported by the Fonds de Recherche du Québec en Santé [scholarship for Master’s Training to A.B., chercheur de mérite award [#295619 to G.B.]; Genome Canada, Genomic Technology Platform [# 245582 to Canadian Center for Computational Genomics and G.B.]; the Canada Research Chair in Transcriptional Genomics [#950-231582 to S.B.]; and the Natural Sciences and Engineering Research Council of Canada [#2019-06490 to S.B.]. Funding for open access charge: Genome Canada, Genomic Technology Platform grant to the Canadian Center for Computational Genomics.

## CONFLICT OF INTEREST

The authors report no conflict of interest.

## TABLE AND FIGURES LEGENDS

Supplementary table 1. Metadata and accession numbers of the read counts files from the RNA-seq data on ENCODE.

Supplementary table 2. Metadata and accession numbers of the alignment files from the ChIP-seq data on ENCODE.

Supplementary table 3. Metadata and accession numbers of the peak files from the ChIP-seq data on ENCODE.

Supplementary table 4. Metadata and accession number of the chromatin interaction and called TADs files from the Hi-C data on ENCODE.

Supplementary table 5. Number of genes labeled as “Up”, “Stable” and “Down” at each timepoint and consensus.

Supplementary figure 1. Distribution of the mean expression of genes. The “Stable” genes have a slight tendency to have a lesser mean expression level across the time-course, but the distribution is overall similar to that of upregulated (“Up”) and downregulated (“Down”) genes.

Supplementary figure 2. Gene expression changes in the RNA-seq samples and Hi-C reproducibility scores. (A) PCA of the normalized RNA-seq data. The top-left, top-right and bottom right panels show the repartition of replicates in the space produced by the first three principal components (PC1 and PC2, PC1 and PC2, then PC2 and PC3, respectively). The bottom-left panel shows the proportion and cumulative proportion of the variance explained by the first five PCs. (B) Heatmap of the quality control scores given by QuASAR-QC. (C) Heatmap of the replication scores given by GenomeDISCO. Comparaisons inside the same timepoint are under the red lines and comparisons across timepoints are over it. (D) Distributions of the scores given by GenomeDISCO between replicates of the same timepoint (blue) or different timepoints (red). (E) Heatmap of the replication scores given by HiC-Spector. (F) Distributions of the scores given byHiC-Spector.

Supplementary figure 3. Count of the nuclear proteins and TAD boundaries. Number of peaks (for nuclear proteins) and of TAD boundaries found in the data.

Supplementary figure 4. The resampling step limits distance bias. (A) Distribution of distances of the “Down-Up” pairs and reference (“Stable-Stable”) pairs before sub-sampling. (B) Distribution of distances of the “Down-Up” pairs and reference (“Stable-Stable”) pairs after the sub-sampling.

Supplementary figure 5. Complete heatmap of odds ratios for the presence of a physical barrier between the genes of the pairs, for all available TFs. (A) The “Stable_Stable” pairs are used as reference. (B) When using the pairs with genes of same comportment (“Stable_Stable”, “Up_Up” and “Down_Down”) as reference, the results are similar, suggesting the “Stable_Stable” category is not only composed of only unregulated genes and is a valid reference. *P-value < 0.05; ***P-value < 0.001. The p-values are empirical, computed with 1000 resampling events.

Supplementary figure 6. Analyzing gene conformation with eQTLs. Pairs of genes where both genes are affected by the same eQTL are enriched for being on the same strand without barrier (CTCF and Cohesin or a TAD boundary) and depleted for being on different strands and separated by a barrier, as compared to control pairs. Here, the distance between genes is defined as the intergenic space. *P-value < 0.025 or p-value > 0.975; ***P-value < 0.001 or p-value = 1. The p-values are empirical, computed with 1000 resampling events.

## Notes

### Competing Interest Statement

The authors have declared no competing interest.

