## Supplementary figures for "Genes on Different Strands Mark Boundaries Associated with Co-regulation Domains"

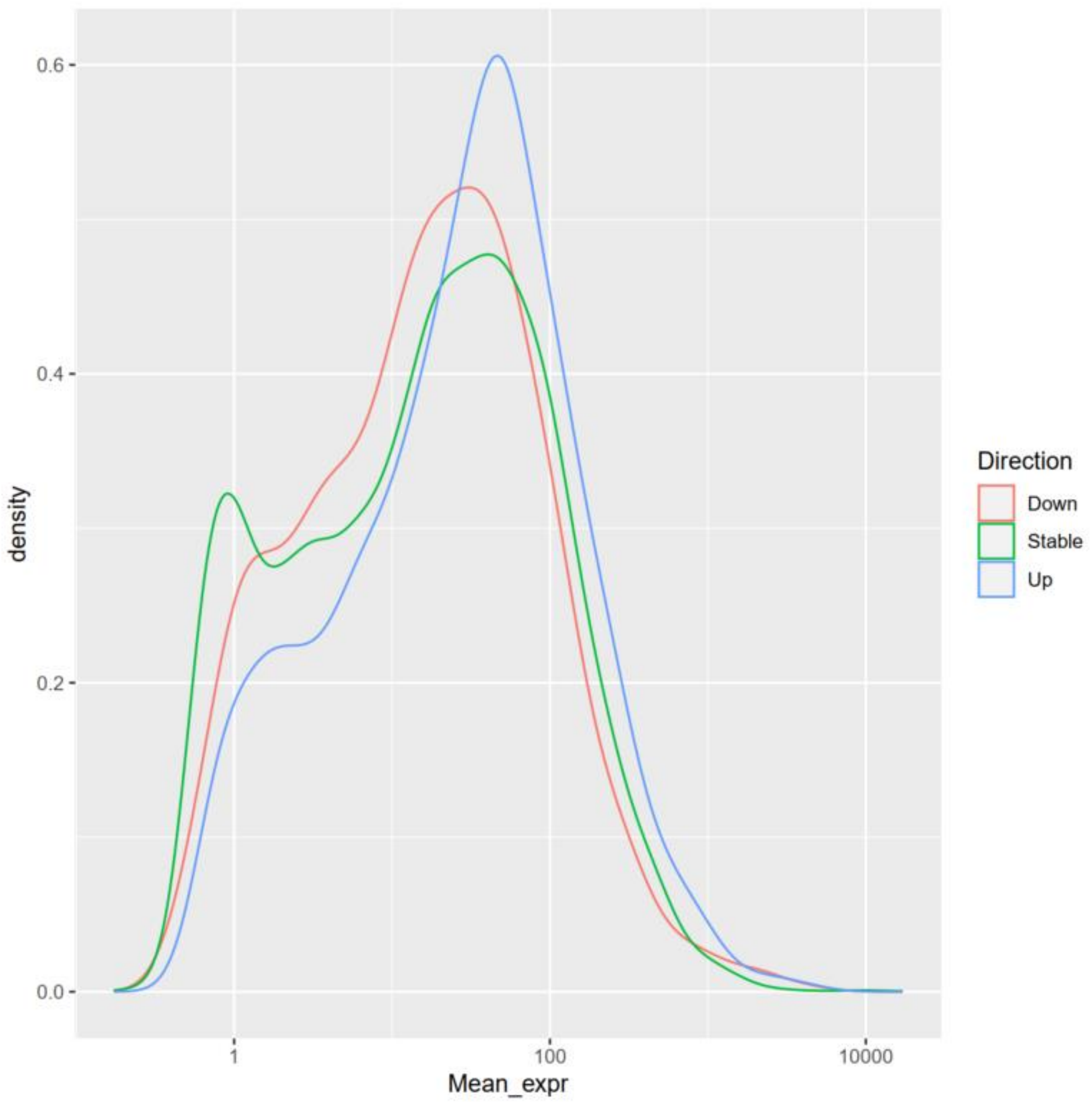


Supplementary figure 1. Distribution of the mean expression of genes. The “Stable” genes have a slight tendency to have a lesser mean expression level across the time-course, but the distribution is overall similar to that of upregulated (“Up”) and downregulated (“Down”) genes.

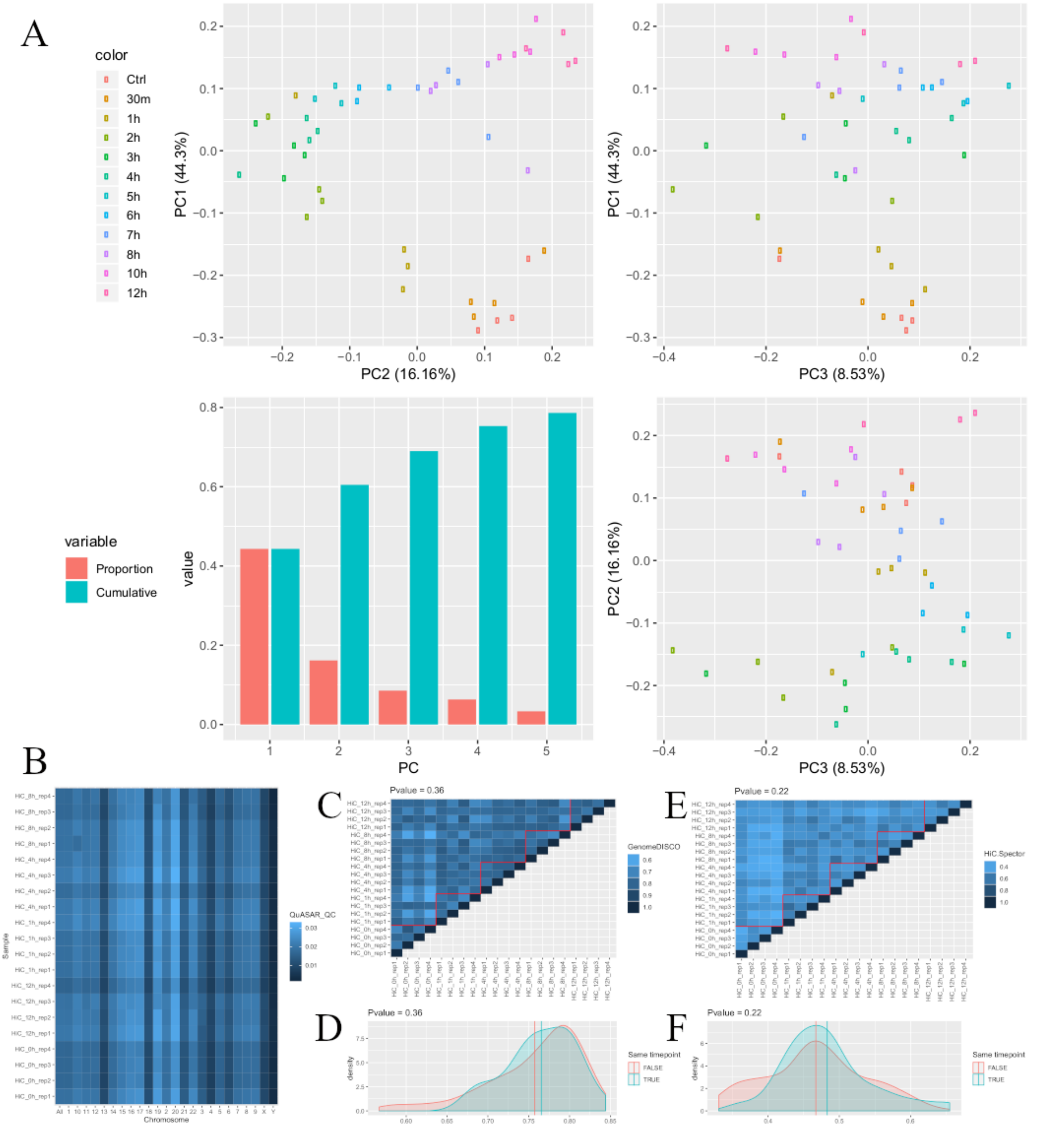


Supplementary figure 2. Gene expression changes in the RNA-seq samples and Hi-C reproducibility scores. (A) PCA of the normalized RNA-seq data. The top-left, top-right and bottom right panels show the repartition of replicates in the space produced by the first three principal components (PC1 and PC2, PC1 and PC2, then PC2 and PC3, respectively). The bottom-left panel shows the proportion and cumulative proportion of the variance explained by the first five PCs. (B) Heatmap of the quality control scores given by QuASAR-QC. (C) Heatmap of the replication scores given by GenomeDISCO. Comparaisons inside the same timepoint are under the red lines and comparisons across timepoints are over it. (D) Distributions of the scores given by GenomeDISCO between replicates of the same timepoint (blue) or different timepoints (red). (E) Heatmap of the replication scores given by HiC-Spector. (F) Distributions of the scores given byHiC-Spector.

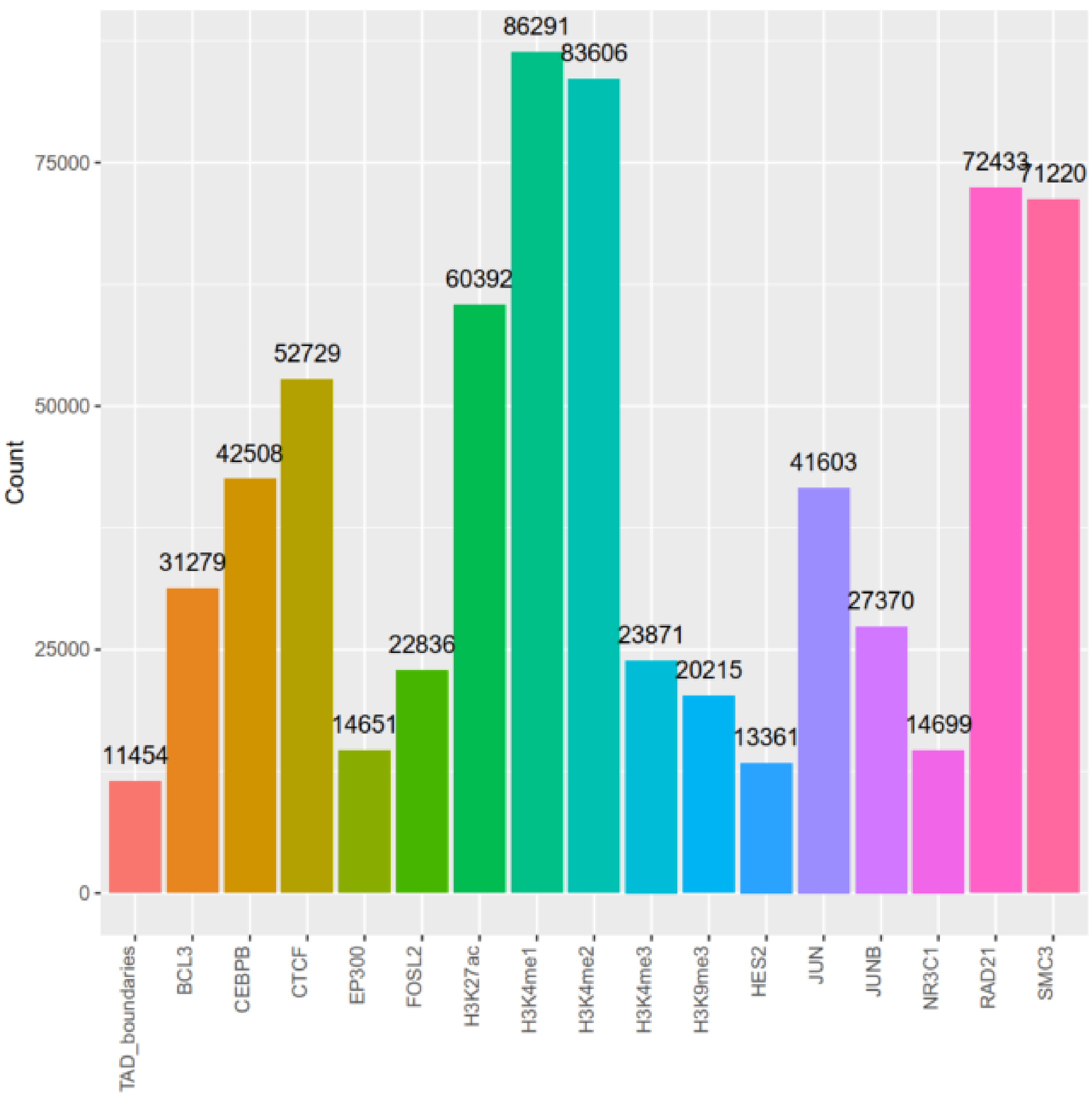


Supplementary figure 3. Count of the nuclear proteins and TAD boundaries. Number of peaks (for nuclear proteins) and of TAD boundaries found in the data.

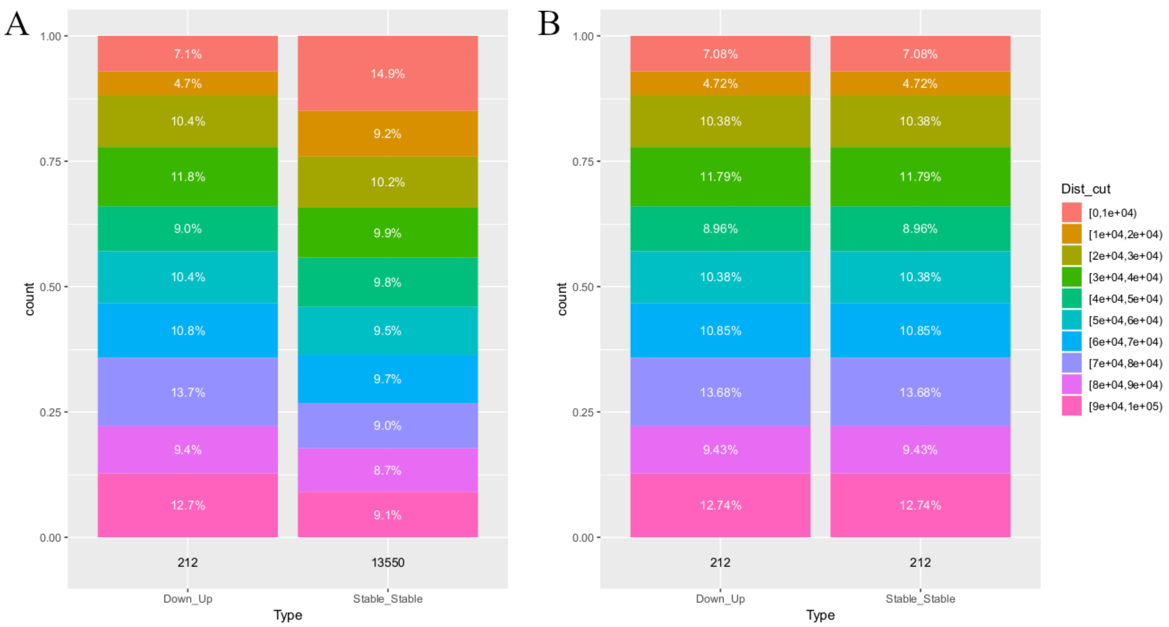


Supplementary figure 4. The resampling step limits distance bias. (A) Distribution of distances of the "Down-Up" pairs and reference ("Stable-Stable") pairs before sub-sampling. (B) Distribution of distances of the "Down-Up" pairs and reference ("Stable-Stable") pairs after the sub-sampling.

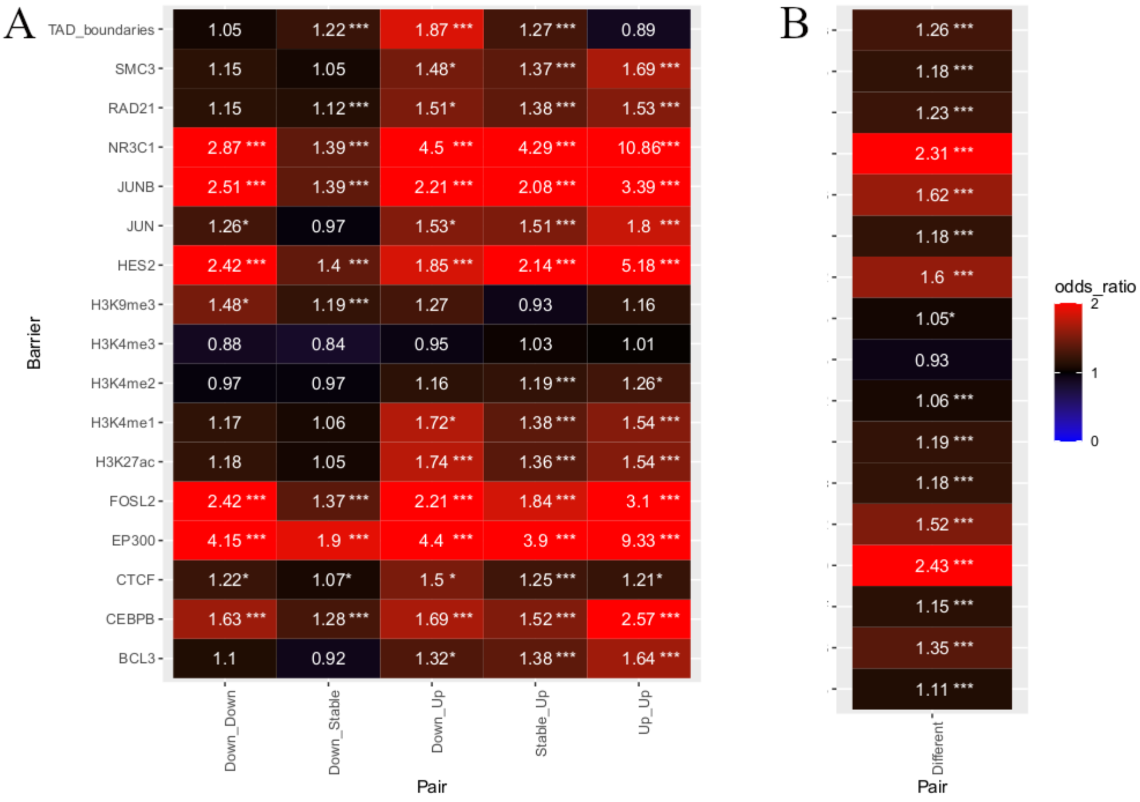


Supplementary figure 5. Complete heatmap of odds ratios for the presence of a physical barrier between the genes of the pairs, for all available TFs. (A) The “Stable_Stable” pairs are used as reference. (B) When using the pairs with genes of same comportment (“Stable_Stable”, “Up_Up” and “Down_Down”) as reference, the results are similar, suggesting the “Stable_Stable” category is not only composed of only unregulated genes and is a valid reference. *P-value < 0.05; ***P-value < 0.001. The p-values are empirical, computed with 1000 resampling events.

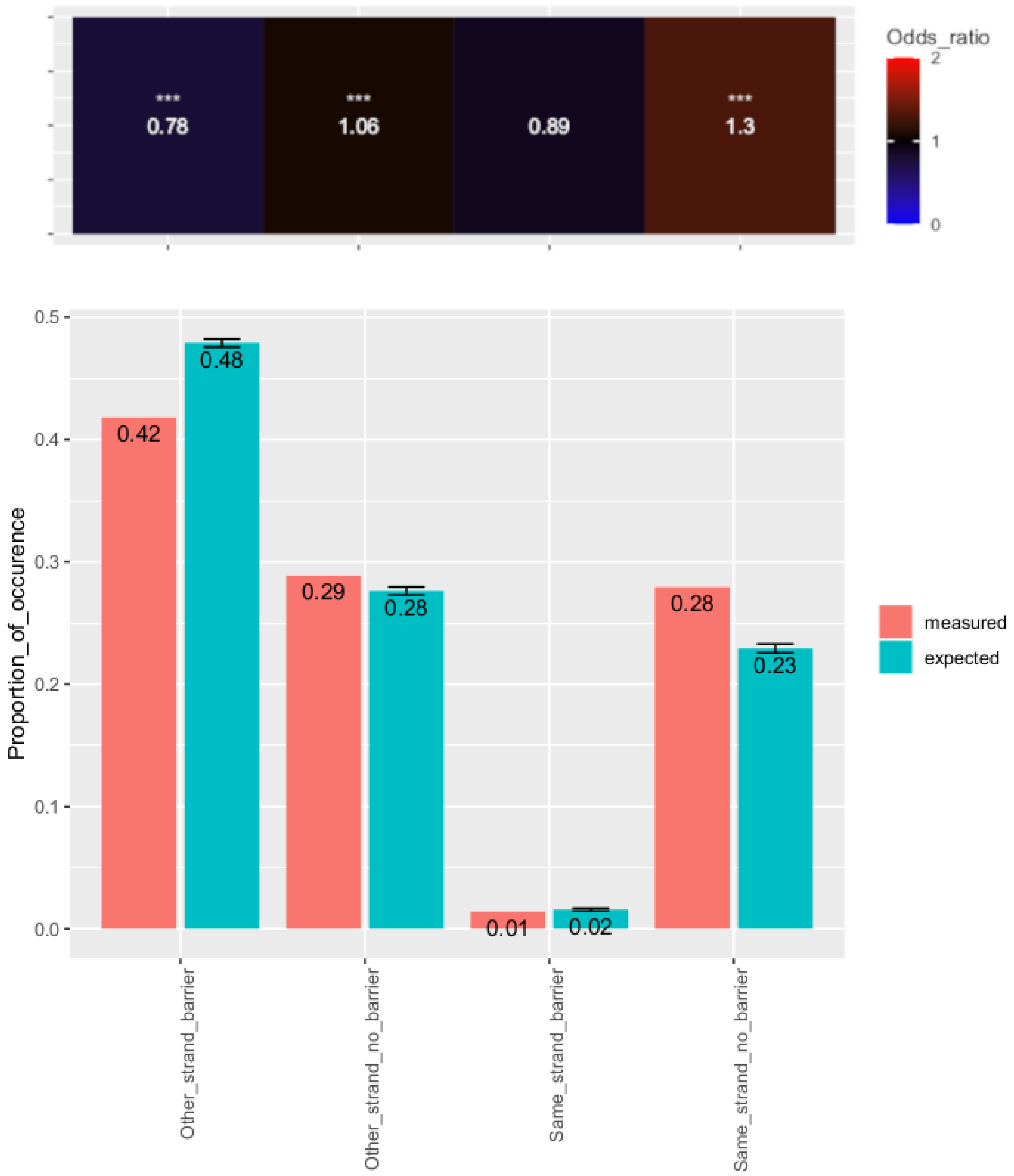


Supplementary figure 6. Analyzing gene conformation with eQTLs. Pairs of genes where both genes are affected by the same eQTL are enriched for being on the same strand without barrier (CTCF and Cohesin or a TAD boundary) and depleted for being on different strands and separated by a barrier, as compared to control pairs. Here, the distance between genes is defined as the intergenic space. *P-value < 0.025 or p-value > 0.975; ***P-value < 0.001 or p-value = 1. The p-values are empirical, computed with 1000 resampling events.
